# Kandinsky: enabling neighbourhood analysis of spatial omics data for functional insights on cell ecosystems

**DOI:** 10.1101/2025.07.10.664141

**Authors:** Pietro Andrei, Mariachiara Grieco, Amelia Acha-Sagredo, Pawan Dhami, Kathy Fung, Manuel Rodriguez-Justo, Matteo Cereda, Francesca D. Ciccarelli

## Abstract

Spatially resolved omics technologies enable investigation of how cells interact within their local environments or neighbourhoods directly *in situ*. Although a few computational methods have been developed to aid this analysis, significant limitations still exist in the way neighbourhoods are defined and exploited for downstream analyses. Here, we present Kandinsky, a computational tool that implements multiple approaches for neighbourhood identification, enabling high flexibility and versatility to address a variety of biological questions. Once identified, Kandinsky applies neighbourhoods for downstream studies, including proximity-based cell grouping for functional comparisons, spatial co-localisation and dispersion, and identification of hot and cold expression areas within the tissue. We apply Kandinsky to transcriptomic and proteomic data from different spatial technologies to showcase how it can reveal functional interactions between cells across multiple biological contexts.

**Availability and implementation:** Kandinsky is freely available as an R package at https://github.com/ciccalab/Kandinsky.

## INTRODUCTION

Recent technological developments in high dimensional spatial biology open new opportunities to study cell properties and interactions *in situ* at unprecedented levels of resolution and throughput. Spatial proteomic approaches such as imaging mass cytometry (IMC)(Giesen, et al., 2014) and CODEX(Goltsev, et al., 2018) (now PhenoCycler(Taube, et al., 2021)) were the first to enable quantification of tens of proteins at micro or nano scale using metal-tagged or fluorescent markers. Imaging-based spatial transcriptomic technologies such as MERSCOPE(Moffitt, et al., 2018), CosMx(He, et al., 2022), or Xenium(Janesick, et al., 2023) simultaneously track the spatial location of RNAs from hundreds or thousands of genes. Sequencing-based approaches such as Slide-Seq(Stickels, et al., 2021), Visium(Stahl, et al., 2016), and Visium-HD(Nagendran, et al., 2023) detect the whole-transcriptome within capture areas or spots of variable size returning different levels of spatial resolutions. Most recent platforms such as COMET(Rivest, et al., 2023) or G4X(Maurer, et al., 2025) even enable quantification of both transcripts and proteins from the same cells.

In parallel with technological advances, several computational tools have been adapted from single cell analysis or developed *ex novo* for spatial data to enable cell segmentation(Jones, et al., 2024; Polanski, et al., 2024), deconvolution(Kleshchevnikov, et al., 2022; Mages, et al., 2023), and phenotyping(Danaher, et al., 2022; Lee, et al., 2025; Singhal, et al., 2024), as well as infer cell-cell communication(Armingol, et al., 2024; Cang, et al., 2023; Zhu, et al., 2024). One of the main advances introduced by spatial approaches is the opportunity to investigate how cells interact *in situ* within the local neighbourhood constituted by their proximal cell environment. Neighbourhood analysis has been used to evaluate cell co-localisation or dispersion of cells within the tissue(Agrawal, et al., 2024; Chen, et al., 2023; Kojima, et al., 2024; Lafzi, et al., 2024; Liu, et al., 2024; Palla, et al., 2022; Varrone, et al., 2024), map gene co-expression patterns in space(Moses, et al., 2023; Withnell and Secrier, 2024), and identify local clusters based on cell composition(Chen, et al., 2023; Ding, et al., 2025; Feng, et al., 2023; Lafzi, et al., 2024; Liu, et al., 2024; Varrone, et al., 2024).

An additional aspect that is currently only partially explored by available analytical approaches is how the local neighbourhood affects the cell phenotype. Some approaches enable functional comparisons between cell types grouped according to the presence or absence of another cell type within the same neighbourhood(Kim, et al., 2023; Mason, et al., 2024), while others restrict this comparisons to gene expression programmes pre-inferred from the data(Agrawal, et al., 2024). However, the flexibility in defining the cell of interests, their neighbours, or the downstream analyses is still limited.

To overcome these limitations we developed Kandinsky, an R package that implements several approaches for cell or spot neighbourhood identification and analysis, including supervised and unsupervised clustering for downstream functional investigations, spatial co-localisation or dispersion, and detection of patterns of high or low gene expression within the tissue. Here, we describe its implementation and provide examples of how Kandinsky can be used to address a variety of scientific questions in different biological contexts.

## METHODS

### Kandinsky data structure and initialization

Kandinsky implements R object-oriented programming defined by an S4 class with 11 components. These components store information regarding c/s-NBs identity, cell/spot polygons, c/s-NB definition, c/s-NBs composition, fixed spot distance (spot data), transcript and FOV coordinates (cell data), texture features of Haematoxylin & Eosin (H&E) staining and tissue slide images.

Class construction is implemented in two steps. First, Kandinsky exploits a set of platform-specific *prepare_seurat* functions which read raw data files from different spatial technologies to initialize a Seurat object. Second, the *kandinsky_init* function creates the object that is stored within the object containing polygon, neighbour and image data associated with the spatial dataset. If a Seurat object has been already created prior to Kandinsky, *kandinsky_init* can still be called if all the required input files are provided. The final output is an updated version of the input Seurat object with the new Kandinsky object stored as a ‘tools’ slot. (**Figure 1A**).

**Figure 1.**
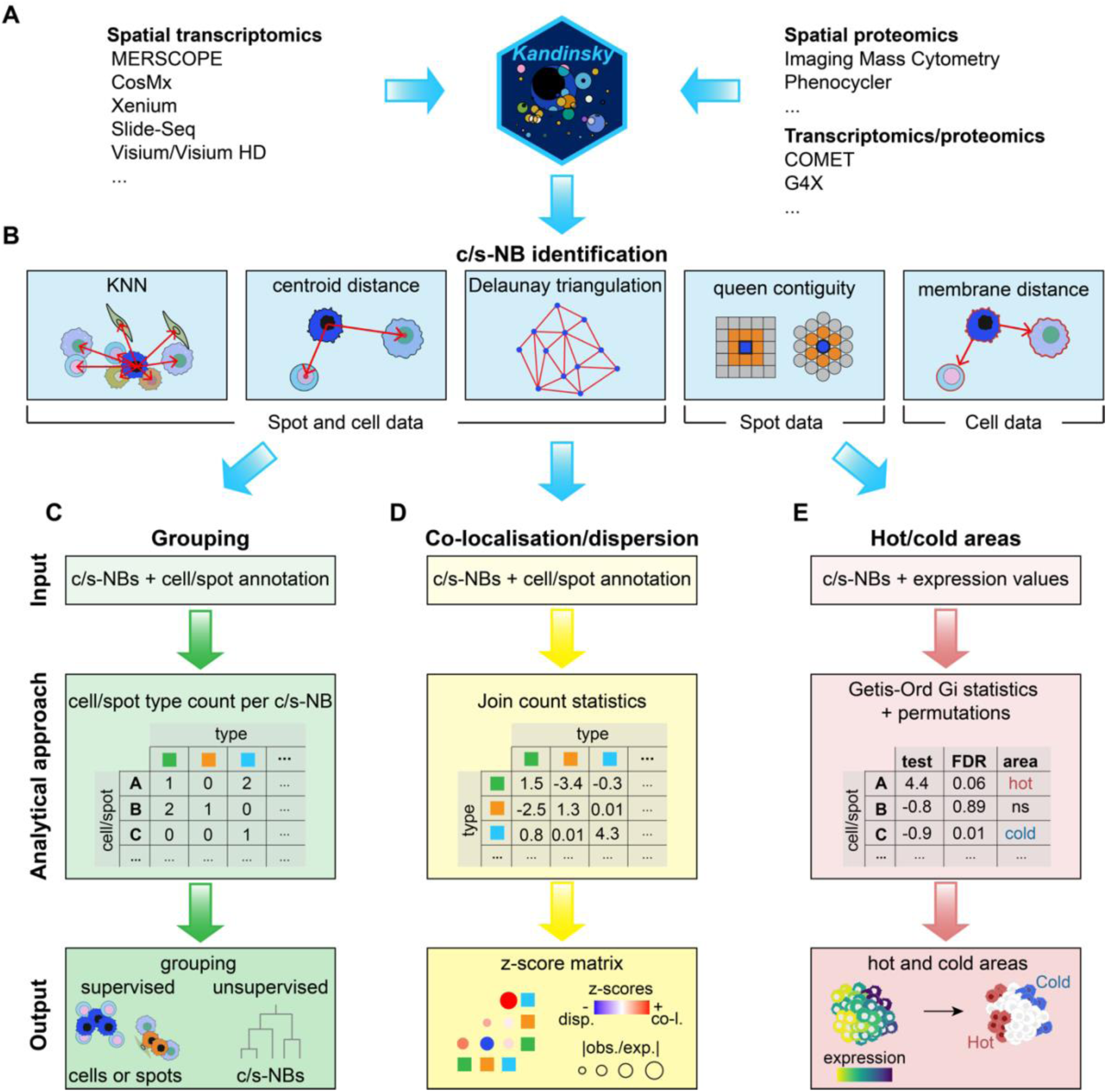
Overview of Kandinsky. A. Examples of technologies delivering spatially resolved data that can be used as an input for Kandinsky. **B.** Representation of the approaches implemented in Kandinsky to identify c/s-NBs based on spatial relationships between cells or spots. KNN, centroid distance, and Delaunay triangulation can be applied to any data type. Queen contiguity and membrane distance can be used only with spot or cell data, respectively. **C.** Cell, spot or c/s-NB grouping. Cell or spot types are labelled according to user-defined annotations, and a matrix is derived by counting the number of cells or spots of every type within each c/s-NB. Cells and spots are grouped based on user-defined neighbourhood criteria (supervised approach). c/s-NBs are grouped based on their cell/spot composition (unsupervised approach). **D.** Cell/spot co-localisation and dispersion. c/s-NBs are used to estimate the tendency of cell or spot types to occupy proximal or distant positions based on z-scores from multi-class join count test. **E.** Hot and cold expression areas. Getis-Ord Gi statistics and permutations are used to identify areas of significantly high and low expression values of a gene, protein, or signature of interest across cells or spots within each c/s-NB. c/s-NB, cell/spot neighbourhood; KNN, K-nearest neighbour.

The *kandinsky_init* function uses cell or spot coordinates to derive cell or spot polygons and identify c/s-NBs relying on R packages sf(Pebesma, 2018), spdep(Bivand, 2018) and tripack(Gebhardt, 2024) based on five neighbouring methods (**Figure 1B**):

1. KNN (based on spdep *knearneigh* function), whereby neighbouring cells are selected as the top user-defined k nearest cells;
2. centroid distance (based on spdep *dnearneigh* function), whereby cells or spots with centroids closer than a user-defined distance are considered within the same c/s-NB;
3. Delaunay triangulation (based on tripack *tri.mesh* function), whereby c/s-NBs are the edges of a network of triangle whose nodes are cells or spots;
4. queen contiguity (spot data only, based on spdep *poly2nb* function), whereby s-NBs are defined as the top user-defined number of layers of surrounding spots;
5. membrane distance (cell data only), whereby cells with membranes closer than a user-defined distance are considered within the same c-NB. Membrane distances are measured by expanding single-cell boundaries using the *st_buffer* function of a radius *r* calculated as

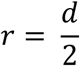

where 𝑑 is the user-defined distance.

Neighbours of each cell are identified by checking the intersection between the expanded single-cell boundaries using the sf *st_intersects* function.

For Visium and Visium HD technology, H&E images are loaded as raster objects using the R package terra(Hijmans, 2025) and aligned to spot coordinates. The resulting list of polygon, c/s-NB, and image objects are assembled into the class as slots, stored as a tool slot within the previously initialised Seurat object and used as input for downstream analysis.

### Cell, spot, and c/s-NB grouping

Input for this analysis are c/s-NBs and an external annotation of cell or spot types used by the *nnMat* function to generate a c/s-NB composition matrix reporting the count of cell or spot types neighbouring each cell or spot across all c/s-NBs (**Figure 1C**). This matrix is then used to derive cell, spot or c/s-NB groups.

For cells and spots, the *nn_query* function is used to derive groups based on user-defined criteria (supervised approach) and perform an optional differential gene expression analysis via the internal call of the *FindMarkers* function.

For c/s-NBs, the *nbCluster* function performs c/s-NB grouping via unsupervised clustering based on composition similarity using the R package mclust(Scrucca, 2023). Users can specify different configurations for the number of clusters to test. For each configuration, the Bayesian Information Criterion (BIC) is evaluated and the cluster configuration with the lowest BIC value is chosen as optimal. Groups will be stored in the metadata and in the sf data slot of the object.

### Analysis of cell or spot co-localisation and dispersion

Input for this analysis are c/s-NBs and an external annotation of cell or spot types used by the *jc_coloc* function that implements the multi-class join count statistics(SOKAL and ODEN, 1978) as defined in the joincount.multi function from the *spdep* package(Bivand, 2018) (**Figure 1D**). The function calculates the number of observed and expected neighbours between each cell or spot type pairs (join counts, *J*) given the overall neighbour composition and under the null hypothesis of random distribution of cell or spot types in the tissue. Then it evaluates if each join occurs by chance by measuring a z-score *z* as:

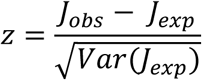

where 𝐽_𝑜𝑏𝑠_ and 𝐽_𝑒𝑥𝑝_ is the observed and expected number of join counts, respectively. Positive z-score values indicate spatial co-localisation. Negative z-score values indicate spatial dispersion. Additionally, the observed-to-expected ratio *r* of join counts is calculated as:

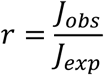

The numerical values of z-scores *z* and their ratio are stored in a tabular format and visualised as a heatmap.

### Hot/cold expression areas

Input for this analysis are c/s-NBs and gene, protein, or gene signature expression values used by the *hotspot_analysis* function to classify cells or spots based on Getis-Ord Gi statistics calculated for each cell or spot using the *localG_perm* function of the spdep package(Bivand, 2018) (**Figure 1E**). Getis-Ord Gi statistics estimates the tendency of cells or spots within the same c/s-NB to express a gene, protein, or signature at significantly higher or lower values than the other cells or spots given the overall c/s-NB composition(Ord, 1995). For each cell or spot 𝑖, the Getis-Ord Gi statistics is calculated as:

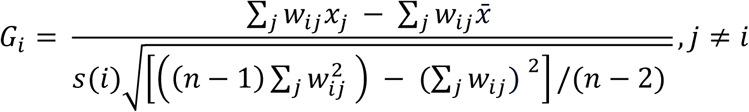

given:

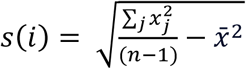

where 𝑤_𝑖𝑗_ is the spatial weight between the cell or spot of interest *i* and any other cell spot *j* within the corresponding c/s-NBs, 𝑥_𝑗_ is the expression value of the variable of interest in any cell or spot 𝑗 within the c/s-NBs, x̄ is the mean expression of the variable of interest across all cells or spots, *s* is the standardized difference between observed and expected expression values across neighbours, and *n* is the total number of cells or spots considered.

Statistical significance is assessed by comparing each observed value to the expected one calculated after randomly reshuffling expression values across all cells or spots via permutations test. An empirical p-value is calculated using the spdep *localG_perm* function for each cell or spot as:

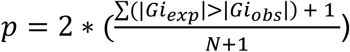

where *Gi*_exp_ and *Gi_obs_* are the expected and observed Getis-Ord Gi statistics, respectively, and *N* is the number of permutations. Empirical p-values are corrected for multiple testing using Benjamini-Hochberg. Cells or spots associated with a positive or negative Getis-Ord Gi statistics and an FDR lower than a user-defined threshold are classified as hot and cold areas, respectively.

### Additional functions

In addition to neighbourhood-based analysis, Kandinsky implements additional function to facilitate spatial data analysis. These include:

1. *get_visium_textures* function to extract texture features from H&E images associated with Visium and Visium HD data. The R package GLCMTextures(Ilich, 2025) is internally used to calculate grey-level co-occurrence matrix texture metrics across pixels. Pixel metrics are then aggregated at the spot level. The output is a spot-metrics matrix stored as slot of the Kandinsky object, with each metrics summarised as mean pixel-level values observed for each spot;
2. *he_mask* function to perform automatic tissue masking from H&E images given a pixel filtering threshold specified via ‘sd_thresh’ paremeter;
3. *global_univ_spatcor* and *global_biv_spatcor* functions to calculate univariate and bivariate spatial correlation coefficients, respectively, for any list of genes or proteins specified by the user;
4. *stitch_samples* function to merge independent samples into a regular grid and create a common coordinate system.

### CosMx normal pancreas dataset

CosMx Human Whole Transcriptome dataset and annotation file of a human pancreas sample was downloaded from the NanoString website(Biology, 2024). The *prepare_cosmx_seur*at and *kandinsky_init* functions were used to initialize a Seraut object with CosMx raw data. c-NBs were defined during initialization using the membrane distance method (‘nb.method=M’ and ‘d.max’ = 0). The *nnMat* and *nn_query* functions were used to (1) build the c-NB composition matrix with CosMx cell type annotation as reference and (2) separate acinar cells into peri-islet and non peri-islet cells based on the presence or absence of islet cells within the same c-NB. Expression profiles of the two acinar cell groups were compared via differential gene expression analysis using *FindMarkers* directly called via the *nn_query* function. Differentially expressed genes were defined as fold change >1.1 or <0.9 and FDR <0.01 (Bonferroni correction). Genes expressed in <20% of peri-islet acinar cells and <5% of non peri-islet acinar cells were excluded to avoid possible contamination.

### IMC PDAC dataset

A Seurat object containing all cell measurements and metadata for all IMC regions of interest (ROIs) was downloaded from the Zenodo website(Sussman, et al., 2024). Cell centroid coordinates from independent ROIs were shifted and merged into a unique coordinate system using the *stitch_samples* function. Following the original methods(Sussman, et al., 2024), a Seurat object was initialized with the *kandinsky_init* function and c-NBs were defined using KNN method (‘nb.method=K’ and ‘k’=20). The *nbCluster* function was then used to group c-NBs across samples, setting the expected number of groups as in the original publication(Sussman, et al., 2024) (‘n_clust’ = 10). Enrichment of PDAC cell types or spatial niches in c-NB groups was tested using Fisher’s test corrected for multiple testing with Benjamini-Hochberg correction.

### Xenium human breast cancer dataset

Xenium Prime 5K Human Pan Tissue and Pathways Panel data of a human breast cancer sample were downloaded from 10X website(Genomics, 2024). Raw Xenium data were loaded into R and used to create a Seurat object using the *prepare_xenium_seurat* function. Cells with <25 transcripts and <20 unique genes were excluded. A new Seurat object was initialized with the *kandinsky_init* function and c-NBs were defined using the cell centroid distance method (‘nb.method=C’ and ‘d.max’=40). For cell clustering, gene expression values were normalized with the *NormalizeData* function with default parameters, and the top 2000 variable genes were selected using the *FindVariableFeatures* function with default parameters. Based on these genes, the *SketchData* function was applied to select a subset of 75000 cells representative of the whole dataset. The top 2000 variable genes were re-selected within the sketched dataset. The *ScaleData* and *RunPCA* functions with default parameters were used to perform dimensionality reduction on the sketched dataset and the *FindNeighbors* and *FindClusters* functions (‘dims’ = 1:50 and ‘res’ = 1.2) were used to identify cell clusters. Uniform manifold approximation and projection (UMAP) was created using the *RunUMAP* function (‘dims’ = 1:50, ‘return.model’ = T). Single cells excluded from sketching were mapped onto the sketched dimensional reduction results using the *ProjectData* function. Cell type annotation for each cluster was manually curated based on the top differentially expressed genes (FC >1.5 and FDR <0.05) identified using the *FindAllMarkers* function.

Co-localisation and dispersion of cell types in the tissue were assessed via multi-class join count statistics using the *jc_coloc* function. Differential gene expression analysis between tumour 5 and 6 cells and the rest of tumour cells was performed using Seurat *FindMarkers* function for all genes expressed in ≥1% of tumour 5 and 6 cells with default parameters. Obtained log2 FC values were used to perform ranked gene set enrichment analysis (GSEA) using the R package fgsea v1.28.0(Sergushichev, 2016).

### CosMx human CRC dataset

CosMx raw data and cell annotations of CRC sample CR48 were downloaded from the Zenodo website(Acha-Sagredo, et al., 2025). Raw files were loaded into R and used to create a Seurat object with the *prepare_cosmx_seurat* function. A new Seurat object was initialized with the *kandinsky_init* function and c-NBs were defined using the cell membrane distance distance (‘nb.method=”M”’ and ‘d.max’=30). Gene expression levels were normalized using *NormalizeData* function.

Hot and cold areas for *CD74* gene expression were derived using the *hotspot_analysis* function to measure the Getis-Ord Gi statistics and statistical significance with 999 permutations (‘perm’ = 999 and ‘padj.thresh’ = 0.05). Enrichment of TAMs, CRC and T/NK cells within *CD74* hot and cold areas was tested using one-tailed Fisher’s exact test.

To assess the effect of T/NK cell proximity on *CD74* expression in CRC cells, the *nnMat and nn_query* functions were run to define the contribution of CRC cell types within each c-NB and group CRC cells based on the presence or absence of T/NK cells within the c-NBs. *CD74* expression between CRC cells proximal to T/NK cells and the rest was compared using two-tailed Wilcoxon’s rank-sum test.

### Visium CRC data generation and analysis

A 5um section was cut from the FFPE block of CRC sample CR48 and placed within the 6x6mm^2^ fiducial frame of the Visium slide. The slide was then incubated at 42 °C for 3h, deparaffinized, H&E stained and imaged using an Olympus VS200 slide scanner. Once imaged, the coverslip was removed, and the slide was decrosslinked. Visium Human Transcriptome Probe kit (v1, PN-1000363) was used for transcript hybridisation. Hybridised RNA molecules were released after tissue permeabilization and captured within each spot by barcoded oligonucleotides. Captured RNA molecules were used for sequencing library preparation and sequenced using NextSeq2000 with a sequencing depth of ≥25000 reads per tissue covered spot.

Visium FASTQ files were processed using the 10X Genomics spaceranger v2.0 software to derive raw gene expression count data. Visium gene expression data were imported in R using Seurat(Hao, et al., 2024). Spots matching with empty regions within the tissue slides annotated using the 10X Genomics Loupe Browser v6.2.0 software and those with <500 transcripts and 300 unique genes were excluded. Genes expressed in less than 10 Visium spots were also removed. Visium raw counts were normalized using the *NormalizeData* function. A new Seurat object was initialized with the *kandinsky_init* function. s-NBs were defined using the queen contiguity method (‘nb.method=”Q”’) and the *nb_expand* function to add the second closest spot layer.

Gene expression scores for the seven immune gene signatures (M0-27, CXCL10/TAM-16, CD209/TAM-13, T-8, CD4-17, Cytotoxic-23, NK-33)(Acha-Sagredo, et al., 2025) were calculated and smoothed across spots using the UCell(Andreatta and Carmona, 2021) *AddModuleScore_UCell* (‘maxRank=5000’) and *SmoothKNN* (‘k’ = 20) functions, respectively. Hot and cold areas for the immune signatures and *CD74* gene were defined using the *hotspot_analysis* function as described above. Enrichment of immune signatures within *CD74* hot and cold areas was tested via one-tailed Fisher exact test and corrected for multiple testing with Benjamini-Hochberg method.

## RESULTS

### Kandinsky overview

Kandinsky is an R package for neighbourhood analysis of spatial omics data. Compared to available approaches, it shows higher versatility in terms of type and resolution of input data, neighbour definition, and downstream analyses (**Table S1**).

Kandinsky uses cell or spot coordinates derived from any spatial transcriptomic or proteomic platform and implements helper functions to automate their loading and formatting into a Seurat object(Hao, et al., 2024) (**Figure 1A**, **Methods**). It then defines cell or spot neighbourhoods (c/s-NBs) according to the spatial relationships between cells or spots as inferred with five methods: k-nearest neighbours (KNN), centroid distance, Delaunay triangulation, queen contiguity, and membrane distance (**Figure 1B**). KNN, centroid distance and Delaunay triangulation are applicable to both cell and spot data. Queen contiguity and membrane distance are limited to spot and cell data, respectively. Once defined, c/s-NBs can be used together with cell or spot type annotation and expression values to (a) group cells, spots, or c/s-NBs based on user-defined similarity or proximity criteria (**Figure 1C**), (b) measure cell or spot co-localisation or dispersion in space (**Figure 1D**) and (c) derive hot and cold expression areas within the tissue (**Figure 1E**).

Cells, spots, or c/s-NBs can be grouped based on a c/s-NB composition matrix derived by counting the number of cell or spot types neighbouring each cell or spot within the tissue (**Figure 1C**). Cells or spots are then grouped following a user-defined supervised approach based on their neighbours, for example grouping epithelial cells surrounded only by other epithelial cells or immune cells proximal to a certain type of stromal cells. c/s-NBs are instead grouped with an unsupervised approach based on the similarity of their cell or spot composition. Once defined, groups of cells, spots, or c/s-NBs can be compared in terms of their functional features, for example global transcriptional profiles or expression levels of specific genes, proteins or signatures of interest. Additionally, their occurrence can be quantified and compared across samples or biological conditions.

Spatial co-localisation and dispersion measure the tendency of cells or spots of a given type to co-localise with or disperse from other cell or spot types (**Figure 1D**). In this case, c/s-NBs are used to measure co-occurrence or dispersion between pairs of neighbouring cell or spot types using z-scores and associated observed-to-expected ratios derived using the multi-class join count statistics(SOKAL and ODEN, 1978) (**Methods**).

Lastly, hot and cold areas identify regions of the tissues where genes, proteins or signatures of interest are expressed more or less than expected by chance (**Figure 1E**). Here, c/s-NBs are used to detect areas of high or low expression levels across neighbouring cells using the Getis-Ord Gi statistics(Ord, 1995) and statistical significance is assessed with permutations (**Methods**).

In addition to these three c/s-NB based analyses, Kandinsky implements additional functions to facilitate the analysis of spatial data, including extraction and masking of image texture features, global spatial correlation tests, and coordinate stitching across independent samples.

### Neighbours modify the phenotypes of healthy and pancreatic cancer cells

We applied Kandinsky to investigate how the local neighbourhood affects cell phenotypes in normal and cancerous human pancreas.

As a first case study, we analysed a CosMx single-cell whole-transcriptome dataset composed of 48951 normal pancreatic cells from a healthy donor(Biology, 2024). Based on the original annotation(Biology, 2024), we identified 12 cell types corresponding to major exocrine (acinar and ductal) and endocrine (islet) pancreatic cells, as well as stellate cells and macrophages (**Figure 2A**). Experiments in mouse showed that acinar cells proximal to islets upregulate trypsin and other digestive enzymes(Egozi, et al., 2020) and undergo acinar-to-beta cell reprogramming(Dahiya, et al., 2024). We used Kandinsky to check whether acinar cells located within peri-islet neighbourhoods of human pancreatic tissues exhibit the same phenotype. First, we identified c-NBs composed of cells in physical contact with each other (membrane distance = 0, **Figure 1B**). We then separated acinar cells in two groups: those in physical contact with at least one islet cell (532 peri-islet acinar cells) and those not in contact with islets (33465 non peri-islet acinar cells, **Figure 2B**). Comparing the gene expression profiles between these two groups of acinar cells, we found 15 differentially expressed genes (**Figure 2C**). Among these, digestion-associated genes, such as *CTRB1/2*, *CPA1, CEL, PNLIPRP1*, were significantly overexpressed in peri-islet acinar cells (**Figure 2C**). These cells also showed significantly lower expression of acinar markers (*REG1A* and *REG1B*) and upregulation of insulin-sensitive (*PDIA4*, *MT2A* and *MT1H/X*) and pancreatic progenitor (*GP2*) genes (**Figure 2C**) supporting their tendency to be reprogrammed into beta-like cells(Dahiya, et al., 2024). Therefore, direct *in situ* human tissue analysis confirmed islet-induced phenotypic changes in acinar cells.

**Figure 2.**
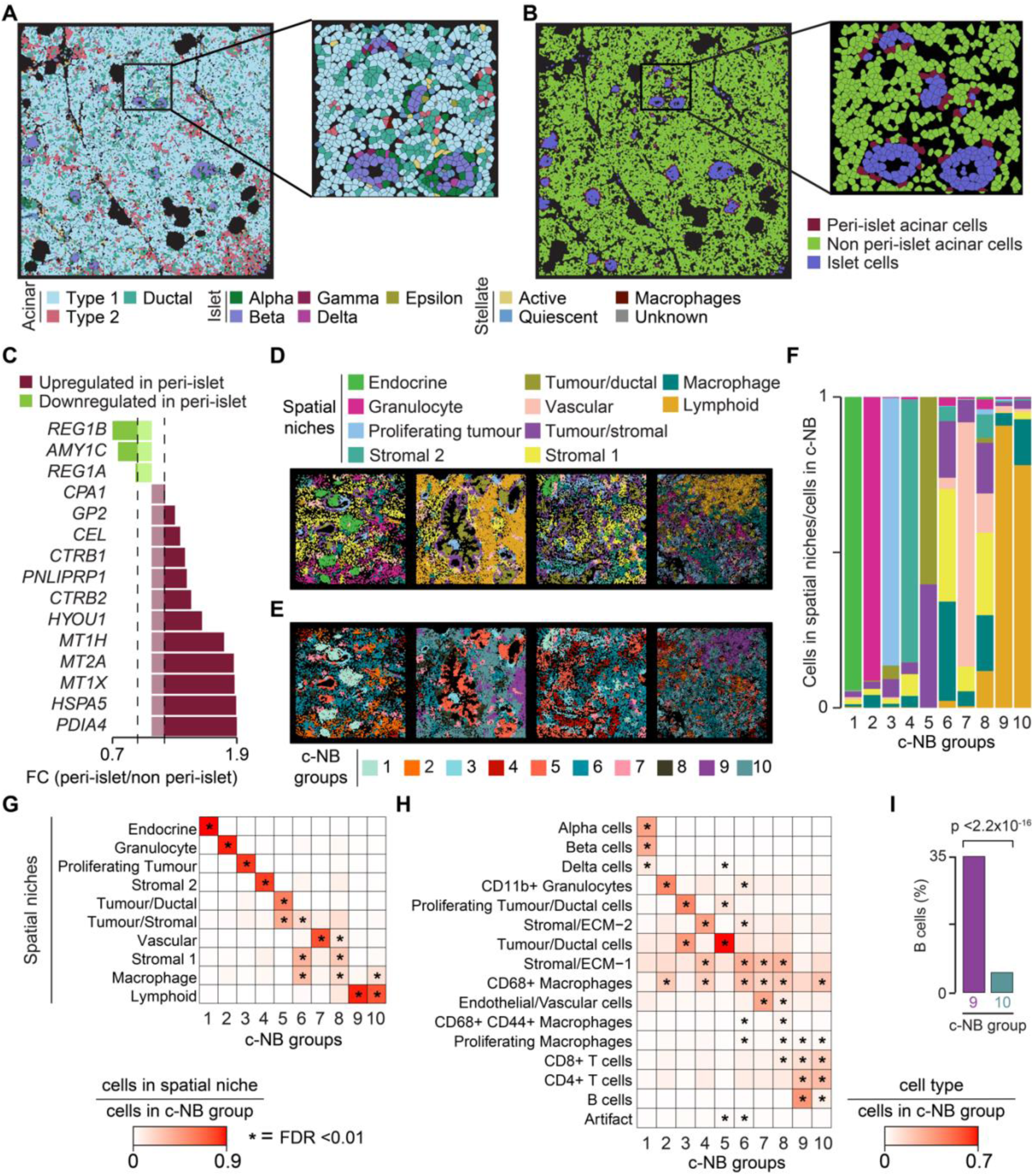
Neighbour-based phenotyping of pancreatic cells. A,B. Cells from nine representative FOVs of a human pancreas sample(Biology, 2024) coloured using the original cell type (**A**) and peri-islet and non peri-islet acinar cell definition from Kandinsky (**B**). Inlets show examples of pancreatic islets surrounded by acinar cells. **C.** Genes up-(FC >1.1) and down-(FC <0.9) regulated in peri-islet acinar cells. Differential gene expression analysis was run between per-islet and non peri-islet acinar cells. Genes expressed in <20% of peri-islet acinar cells and <5% of non peri-islet acinar cells were excluded to limit transcript contamination. Genes with Bonferroni adjusted p-value <0.01 were considered differentially expressed. **D,E.** Cells from representative FOVs of four PDAC samples (P1-R1, P4-R1, P6-R3, P8-R2) coloured by spatial niches(Sussman, et al., 2024) (**D**) and c-NB groups (**E**). **F.** Proportion of spatial niches(Sussman, et al., 2024) within c-NB groups. **G,H.** Enrichment of spatial niches (**G**) and cell types (**H**) within c-NB groups. Enrichment was tested using one-tailed Fisher’s exact test. P-values were corrected for multiple testing with Benjamini-Hochberg method. **I.** Comparison of B cell proportions between c-NB groups 9 and 10. Proportions were compared using two-sided Fisher’s exact test. c-NB, cell neighbourhood; FC, fold change; FDR, false discovery rate; FOV, field of view; IMC, imaging mass cytometry; PDAC, pancreatic ductal adenocarcinoma.

As a second case study, we analysed an IMC spatial proteomic dataset composed of 144976 single cells from nine pancreatic ductal adenocarcinoma (PDAC) samples assayed with a panel of 26 metal-tagged antibodies(Sussman, et al., 2024). Applying KNN to derive neighbourhoods and k-means for clustering, the original study described ten distinct spatial niches(Sussman, et al., 2024) (**Figure 2D**). We tested whether Kandinsky could detect similar or even better refined cell neighbourhoods. To define c-NBs, we applied KNN with the same neighbourhood size (k=20) of the original study and grouped these into ten groups using the built-in unsupervised model-based clustering function (**Methods**). Overall, we found good concordance between cells assigned to the original spatial niches and newly detected c-NB groups (adjusted Rand index = 0.4, **Figure 2E**). Four niches (endocrine, granulocyte, proliferating tumour, and stromal 2) showed significant overlap with only one c-NB group (**Figures 2F, G**). For the remaining six niches, c-NB groups refined the original annotation. For instance, cells in the original tumour/ductal and tumour/stromal niches were reassigned to c-NB groups 5 and 6 (**Figures 2G**). Enrichment analysis of cell types confirmed that c-NB group 5 was mostly composed of tumour cells, while c-NB group 6 of a variety of stromal and immune cells (**Figures 2H**). Similarly, the vascular niche, which in the original study contained also a significant proportion of stromal cells and macrophages(Sussman, et al., 2024), was split into c-NB groups 7 (composed mostly of endothelial cells with a minor contribution of macrophages and stromal cells) and 8 (mostly macrophages and stomal cells with a minor contribution of endothelial cells, **Figures 2G,H**). Finally, the original lymphoid niche was further split into c-NB groups 9 and 10 (**Figure 2G,H**). Both c-NB groups were enriched in T cells, but group 9 showed a significant overrepresentation of B cells (**Figure 2I**) while group 10 of macrophages (**Figure 2G,H**). In all these examples, a more precise identification of cell populations within separate groups may improve to characterise their functional properties.

### Myoepithelial cells surround non-invasive cancer cells within the breast tissue

We tested Kandinsky to assess the tendency of cells to significantly co-localise or disperse in the tissue (**Figure 1D**) using a single-cell spatial transcriptomic dataset from a stage II-A breast cancer sample profiled with the Xenium Prime 5K Human Panel(Genomics, 2024). After removing low-quality cells (**Methods**), we performed unsupervised clustering of the 468583 remaining cells leading to the identification of 23 cell populations based on differentially expressed marker genes (**Figures 3A, B**).

**Figure 3.**
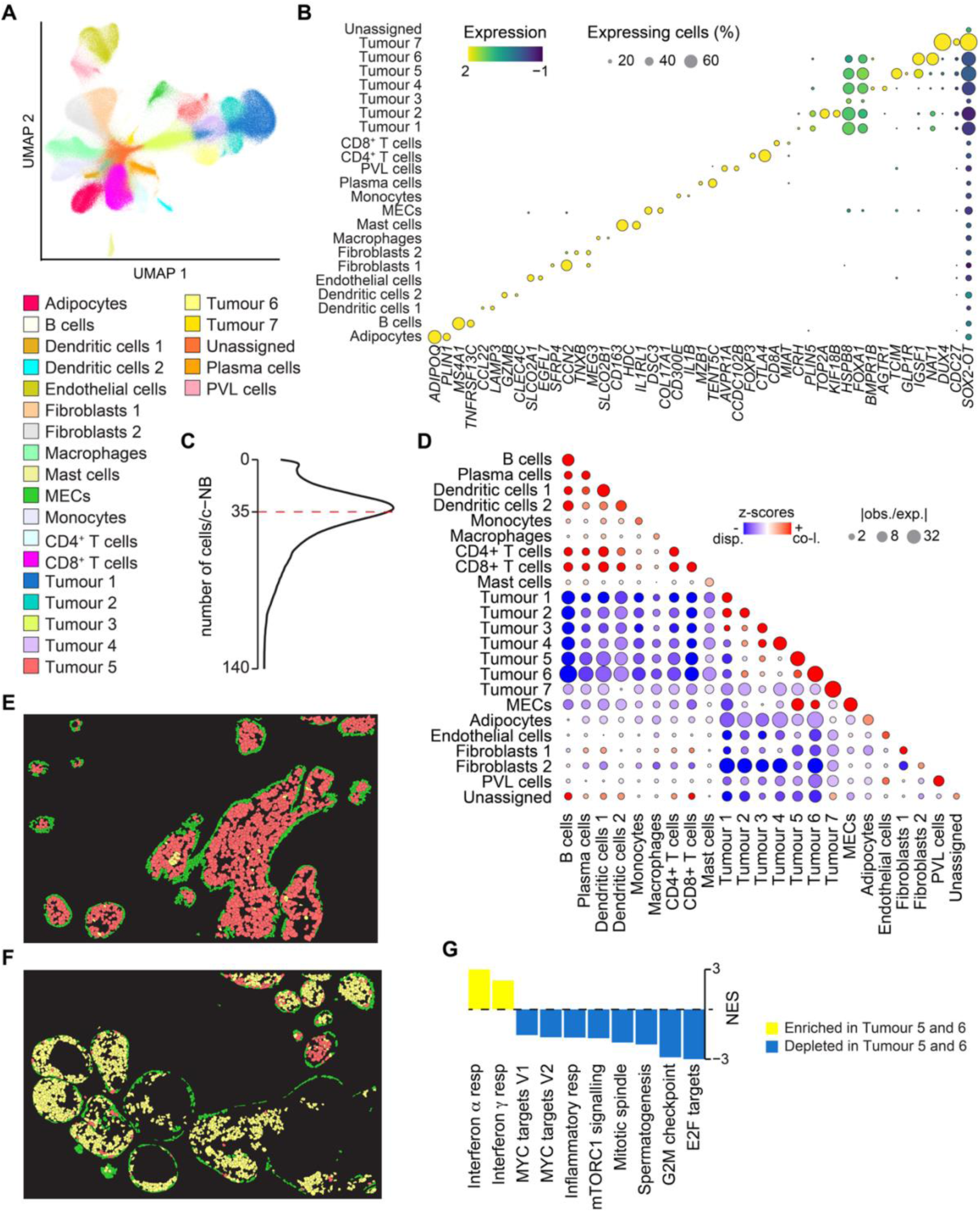
Cell localisation analysis in breast cancer. A. Unsupervised clustering of 468583 cells from a breast cancer single cell transcriptomics dataset obtained with the Xenium 5K technology(Genomics, 2024). Cells are coloured by cell type. **B.** Mean expression value of marker genes representative of cell types identified with clustering. Differential gene expression analysis was performed between each cell type and the rest. Up to two differentially expressed genes per cell type (FC >2, FDR <0.05) were selected for visualisation. **C.** Distribution of cells per c-NB, identified using a centroid distance of 40um. Median number of cells is reported as a dashed red line. **D.** Multi-class join count z-score matrix assessing co-occurrence or dispersion of each cell pair (**Methods**). Positive z-score values indicate co-localisation, negative z-score values indicate dispersion. Dot size indicate observed-to-expected ratio of join counts. **E-F**. Representative regions of the breast tissue sample showing layers of MECs surrounding tumour 5 (**E**) and tumour 6 (**F**) cells. **G.** Significantly enriched (NES >0) and depleted (NES <0) hallmark cancer pathways(Liberzon, et al., 2015) in cancer cells of tumour 5 and 6 clusters compared to the rest of cancer cells. Enrichment was estimated using ranked GSEA. Pathways with FDR <0.05 were considered significant. c-NB, cell neighbourhood; FC, fold change; FDR, false discovery rate (Benjamini-Hochberg); GSEA, gene set enrichment analysis; MECs, myoepithelial cells; NES, normalised enrichment score; PVL, perivascular-like.

We used Kandinsky to define c-NBs according to a centroid distance of 40um (**Figure 1B**), resulting in a median of 35 cells per c-NB (**Figures 3C**). We then quantified the tendency of cells to significantly aggregate or disperse in the tissue using multi-class join count derived z-scores (**Figure 1D**). As expected, immune cell populations tended to localise proximal to each other and to be separated from both tumour cells and, to a lower extent, the stromal compartment (**Figures 3D**). Overall, tumour cells also tended to co-localise in the tissue space, except for a small population corresponding to 1.6% of all tumour cells that clustered away from anything else (tumour 7, **Figure 3D**).

Interestingly, Kandinsky detected spatial co-localisation between myoepithelial cells (MECs) and two tumour clusters (tumour 5 and 6, **Figure 3D**). In normal breast, MECs are known to protect the structure and integrity of epithelial cells(Gudjonsson, et al., 2005; Runswick, et al., 2001). In breast cancer, MECs act as a physical barrier against tumour cell invasion(Adriance, et al., 2005), and the structural integrity of MEC layer is associated with the risk of progression in patients with ductal carcinoma in situ(Risom, et al., 2022; Russell, et al., 2015). When inspecting the spatial organisation of MECs in the tissue, we observed that these cells formed a monolayer around duct-like structures formed of tumour 5 (**Figure 3E**) and tumour 6 (**Figure 3F**) cells. Compared to the other tumour cells, tumour 5 and 6 cells showed higher interferon response and reduced proliferation potential (**Figure 3G**). Since MEC layers surrounding milk ducts are gradually lost during cancer progression as a consequence of the increase in cancer cell proliferation and spreading(Hu, et al., 2008), it is tempting to speculate that these tumour cells might have a limited invasive potential. This analysis shows how studying the spatial organisation of cell populations in the tissue can inform on functional phenotypes.

### CRC tissue areas enriched in lymphocytes and TAMs show high *CD74* expression

We recently showed that cytotoxic lymphocytes induce upregulation of interferon genes, including MHC class II invariant chain *CD74*, in tumour associated macrophages (TAMs) and colorectal cancer (CRC) cells(Acha-Sagredo, et al., 2025). This is clinically relevant because *CD74* is highly expressed in CRC patients that respond to treatment with immune checkpoint inhibitors(Acha-Sagredo, et al., 2025; Bortolomeazzi, et al., 2021; Kim, et al., 2024).

Using Kandinsky, we further analysed whether *CD74* expression levels correlate with the spatial localisation of cancer cells, TAMs and lymphocytes in a CRC sample profiled with the CosMx 1000 gene panel(Acha-Sagredo, et al., 2025). Following the original annotation(Acha-Sagredo, et al., 2025), we found five main cell clusters, corresponding to CRC, T/NK, plasma/B, myeloid, and stroma cells (**Figure 4A**). We then identified c-NBs applying a cell membrane distance of 30um, which resulted in a median of 36 cells per c-NB (**Figure 4B**). Finally, we applied Getis-Ord Gi statistics (**Figure 1E**) on single cell *CD74* expression levels (**Figure 4C**) to map *CD74* hot and cold areas within the sample (**Figure 4D**).

**Figure 4.**
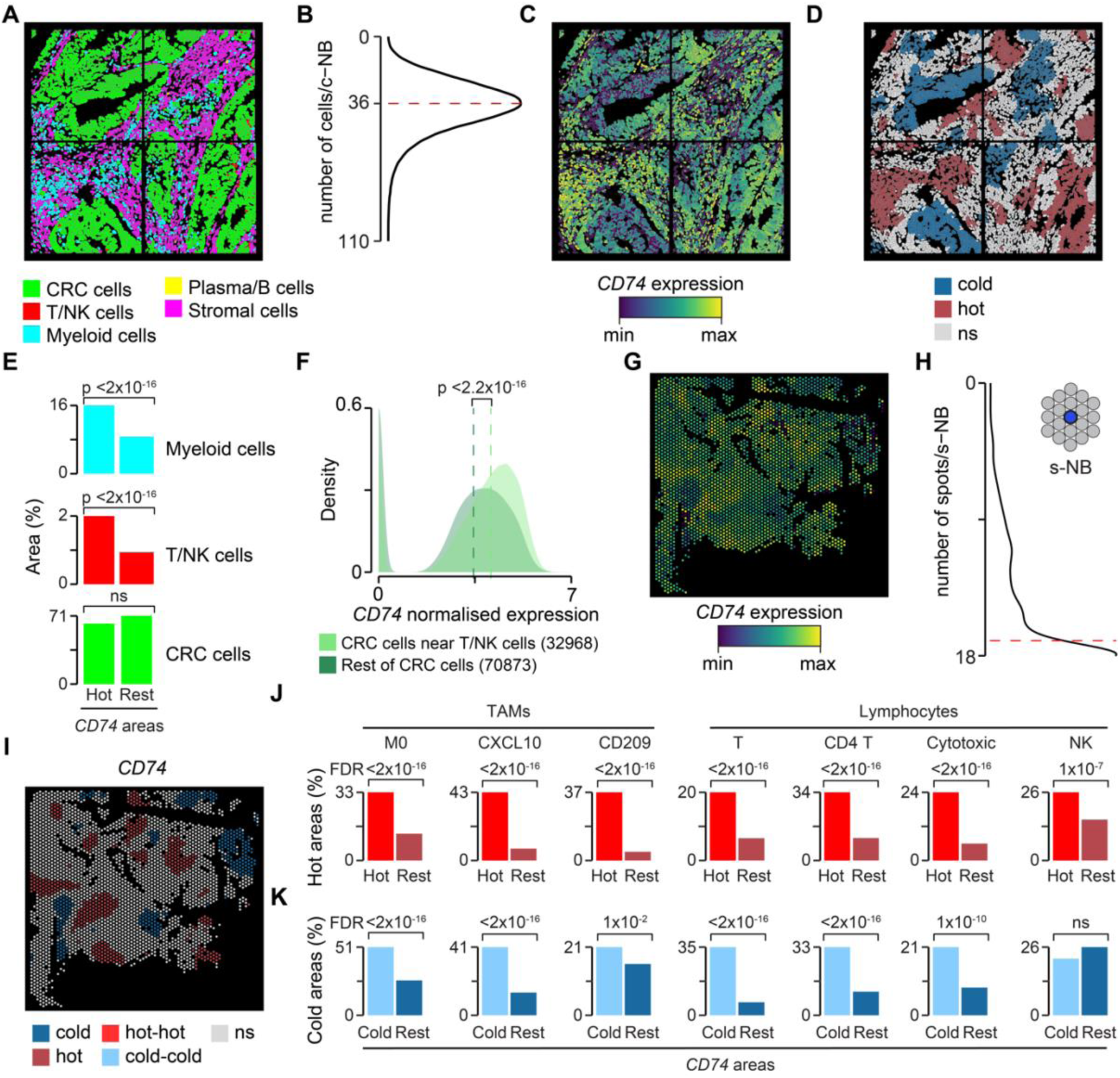
Cell enrichment within *CD74* hot and cold CRC tissue areas. A. Cells from four representative FOVs of a human CRC sample (CR48) coloured by original cell type(Acha-Sagredo, et al., 2025). **B.** Distribution of cells per c-NB, identified using a cell membrane distance of 30um. Median number of cells is reported as a dashed red line. **C,D**. Cells from the FOVs shown in (**A**), coloured by *CD74* expression levels (**C**) and *CD74* hot or cold areas (**D**). *CD74* hot and cold areas were identified using the Getis-Ord Gi statistics applied to neighbour cells. **E.** Comparison of *CD74* hot and non-hot areas containing myeloid, T/NK, or CRC cells. **F.** Density plot of *CD74* expression values in CRC cells with or without T/NK cells within their c-NBs. Distributions were compared using two-sided Wilcoxon’s rank-sum test. **G.** Visium spots for CRC sample (CR48), coloured by *CD74* expression levels. **H.** Distribution of spots per s-NB, identified using the queen contiguity method with the first two contiguous layers surrounding each spot. Median number of spots is reported as a dashed red line. **I.** Visium spots for CRC sample (CR48), coloured by *CD74* hot or cold areas. *CD74* hot and cold areas were identified based on Getis-Ord Gi statistics applied to neighbour spots. **J,K.** Comparison of *CD74* hot and non-hot (**J**) and *CD74* cold and non-cold (**K**) areas overlapping with hot and cold areas for gene signatures associated with TAMs and lymphocyte subpopulations(Acha-Sagredo, et al., 2025). Significance in (D) and (I) was tested based on permutations and cells associated with a positive or negative Getis-Ord Gi statistics with FDR <0.05 were assigned to *CD74* hot or cold areas, respectively (**Methods**). Proportions in (E), (J) and (K) were compared using one-sided Fisher’s exact test and corrected for multiple testing when needed. c-NB, cell neighbourhood; CRC, colorectal cancer; FDR, false discovery rate (Benjamini-Hochberg); FOV, field of view; NK, natural killer; ns, not significant s-NB, spot neighbourhood; TAMs, tumour-associated macrophages.

To test the overlap between spatial localisation of CRC, myeloid and T/NK cells and *CD74* expression levels, we measured the relative enrichment of each cell population in *CD74* hot areas using one-tailed Fisher’s exact test. We found a significant enrichment of myeloid and T/NK cells in *CD74* hot areas (**Figure 4E**), confirming that TAMs are the major source of *CD74* expression in the tumour stroma upon stimulation by interferon-producing lymphocytes(Acha-Sagredo, et al., 2025; Bortolomeazzi, et al., 2021). Unexpectedly, CRC cells showed no significant enrichment in *CD74* hot areas (**Figure 4E**). We reasoned that this may be due to the fact that only CRC cells proximal to lymphocytes would co-localise within *CD74* high expression areas, since only these CRC cells overexpress *CD74* in response to IFN stimulation(Acha-Sagredo, et al., 2025). To test this hypothesis, we compared *CD74* expression between CRC cells in c-NBs also containing T/NK cells and those in c-NBs with no T/NK cells. We confirmed that CRC cells proximal to T/NK cells express significantly higher *CD74* levels than the rest of CRC cells (**Figure 4F**), thus supporting the initial hypothesis.

Being based on 1000 genes only, the CosMx panel enabled broad separation of the main immune populations (**Figure 4A**) but prevented further characterisation of relevant immune subpopulations associated with *CD74* hot and cold areas. To overcome this limitation, we profiled another tissue region of the same CRC sample using Visium whole transcriptomics approach (**Methods**). After quality control and removal of low quality spots and genes, we mapped *CD74* expression levels across the retained 2955 spots (**Figure 4G**). To define s-NBs, we used the queen contiguity approach (**Figure 1B**) considering the first two contiguous layers surrounding each spot, resulting in a median of 17 spots per s-NB (**Figure 4H**).

Like for the c-NBs analysis, we used s-NBs to identify *CD74* hot and cold areas using the Getis-Ord Gi statistics (**Figure 4I**). Since the Visium technology does not provide single cell resolution, we could not assess the enrichment of TAM and lymphocyte sub-populations in *CD74* hot and cold areas directly. Instead, we used seven manually curated gene signatures representing three TAM subpopulations (M0, CXCL10, and CD209 TAMs) and four lymphocyte subpopulations (cytotoxic lymphocytes, T, NK, and CD4 T cells)(Acha-Sagredo, et al., 2025). We identified hot and cold areas in the CRC tissue for each of these gene signatures using again the Getis-Ord Gi statistics and applied one-tailed Fisher’s exact test to assess their overlap with *CD74* hot and cold areas. Overall, we found a significant enrichment of hot areas for signatures associated with TAM and lymphocyte subpopulations within hot areas for *CD74* expression (**Figure 4J**). Similarly, we found areas significantly depleted in TAM and lymphocyte subpopulations within areas with low *CD74* expression (**Figure 4K**). This analysis confirmed *in situ* the *CD74* overexpression in TAM subpopulations induced by T cells previously detected *in vitro*.

## DISCUSSION

Kandinsky is a computational tool designed for cell or spot neighbourhood analysis that allows the user to interact with data derived from several spatial technologies and adapt the analysis to the biological query. Compared to other methods, Kandinsky can identify c/s-NBs with greater flexibility. For example, to our knowledge, Kandinsky is the first tool able to apply single-cell segmentation data to define c-NBs according to cell membrane distance in addition to other commonly implemented criteria (KNN, centroid distance, Delaunay triangulation, queen contiguity). In addition, Kandinsky implements a suite of functions and analytical modules to facilitate spatial data analysis without depending on multiple software for each specific task.

We showed with real-world data how downstream analytical modules implemented in Kandinsky make use of c/s-NBs in a coherent way, applying c/s-NB definition to each analytical task. Users can interrogate c/s-NBs to define group of cells and spots proximal to cell types of interest, or c/s-NBs sharing a similar neighbourhood composition, study spatial co-localisation patterns between cell or spot types, or identify local hot or cold expression areas. By analysing spatial datasets generated through CosMx, IMC, Xenium, and Visium platforms, we demonstrated Kandinsky versatility to produce biologically meaningful results supported by literature evidence.

Although c/s-NB grouping, co-localisation and dispersion analyses are compatible with spatial data of any resolution, they do not support direct inclusion of cell type deconvolution often associated with spot-level data. This is a current limitation of Kandinsky that could be part of future updates. Another possible improvement would be the possibility to run Kandinsky on more than one dataset at the same time for multi-sample comparisons. This requires a substantial change of Kandinsky object format to store molecular and spatial information for independent samples, without losing interoperability with the underlying Seurat infrastructure.

Given the continuous development new of spatial technologies, we aim at maintaining Kandinsky code and documentation up to date and expanding the list of compatible spatial platforms and possible analyses.

## Data Availability

CosMx human pancreas data(Biology, 2024) were downloaded from https://nanostring.com/products/cosmx-spatial-molecular-imager/ffpe-dataset/cosmx-smi-human-pancreas-ffpe-dataset. IMC PDAC data(Sussman, et al., 2024) were downloaded from https://zenodo.org/records/10246315. Xenium breast cancer data(Genomics, 2024) were downloaded from https://www.10xgenomics.com/datasets/xenium-prime-ffpe-human-breast-cancer.

CosMx CRC data(Acha-Sagredo, et al., 2025) were downloaded from https://zenodo.org/records/10927005. Visium CRC data have been deposited in Zenodo (https://doi.org/10.5281/zenodo.15209564). Access to these data is restricted to non-commercial research only and requires data sharing agreement with the Francis Crick Institute.

## Acknowledgements

We thank the Experimental Histopathology and the Advanced Light Microscopy facilities of the Francis Crick Institute for the support with the Visium experiment, and Gabriele Boscagli for reviewing the code.

## Author Contributions

Conceptualization: P.A., and F.D.C.; methodology: P.A., M.G., M.C., and F.D.C.; software: P.A., M.G., and M.C.; formal analysis: P.A.; investigation: A.A.-S., P.D., and K.F; resources: P.D., K.F., and M.R.-J.; writing – original draft: P.A., and F.D.C.; writing – review and editing: P.A., M.G., M.C., and F.D.C.; visualization: P.A., and F.D.C.; supervision: M.C. and F.D.C.; funding acquisition: F.D.C.

